# PartSeg, a Tool for Quantitative Feature Extraction From 3D Microscopy Images for Dummies

**DOI:** 10.1101/2020.07.16.206789

**Authors:** Grzegorz Bokota, Jacek Sroka, Subhadip Basu, Nirmal Das, Paweł Trzaskoma, Yana Yushkevich, Agnieszka Grabowska, Adriana Magalska, Dariusz Plewczyński

## Abstract

**Background:** Bioimaging techniques offer a robust tool for studying molecular pathways and morphological phenotypes of cell populations subjected to various conditions. As modern high resolution 3D microscopy provides access to an ever-increasing amount of high quality images, there arises a need for their analysis in an automated, unbiased and simple way.

Segmentation of structures within cell nucleus, which is the focus of this paper, presents a new layer of complexity in the form of dense packing and significant signal overlap.

At the same time the available segmentation tools provide a steep learning curve for new users with limited technical background. This is especially apparent in bulk processing of image sets, which requires the use of some form of programming notation.

**Results:** In this paper, we present PartSeg, a tool for segmentation and reconstruction of 3D microscopy images, optimised for the study of cell nucleus. PartSeg integrates refined versions of several state-of-the-art algorithms, including a new multi-scale approach for segmentation and quantitative analysis of 3D microscopy images.

The features and user-friendly interface of PartSeg were carefully planned with biologists in mind, based on analysis of multiple use cases and difficulties encountered with other tools, to offer ergonomic interface with a minimal entry barrier. Bulk processing in an ad-hoc manner is possible without the need for programmer support. As the size of datasets of interest grows, such bulk processing solutions become essential for proper statistical analysis of results.

Advanced users can use PartSeg components as a library within Python data processing and visualisation pipelines, for example within Jupyter notebooks. The tool is extensible so that new functionality and algorithms can be added by the use of plugins.

For biologists the utility of PartSeg is presented in several scenarios, showing the quantitative analysis of nuclear structures.

**Conclusions:** In this paper, we have presented PartSeg which is a tool for precise and verifiable segmentation and reconstruction of 3D microscopy images. PartSeg is optimised for cell nucleus analysis and offers multiscale segmentation algorithms best-suited for this task. PartSeg can also be used for bulk processing of multiple images and its components can be reused in other systems or computational experiments.

**Contact**

## 1 Background

For a decade, high-throughput bioimaging techniques offered a robust tool for studying molecular pathways and morphological phenotypes of cell populations subjected to various conditions [1, 2]. Due to recent advances in light and electron microscopy, a large number of input images can be produced in a relatively short time span. Therefore, it becomes critical to extract numerical features from imaging data in an automated, unbiased and simple way.

The cell nucleus is a highly organised and crowded organelle, composed of many functional and structural domains (Fig.1). The past two decades of research show that changes in higher-order chromatin structures, that is, spatial and temporal rearrangements of chromatin, are involved in transcriptional control and other cellular functions [3, 4]. The 3-C based methods, which decipher folding of chromatin, quantify the probability of interaction between two genomic fragments, such as promoters and enhancers, that are in close spatial proximity, but may be dispersed far across the genome. To refine this biochemical data, showing the outcome averaged over millions of cells, precise microscopic analysis of nuclear components like genes, chromatin and nuclear domains is necessary. Therefore, microscopy is often used for verification, at scales of individual cells or even genetic copies.

**Figure 1:**
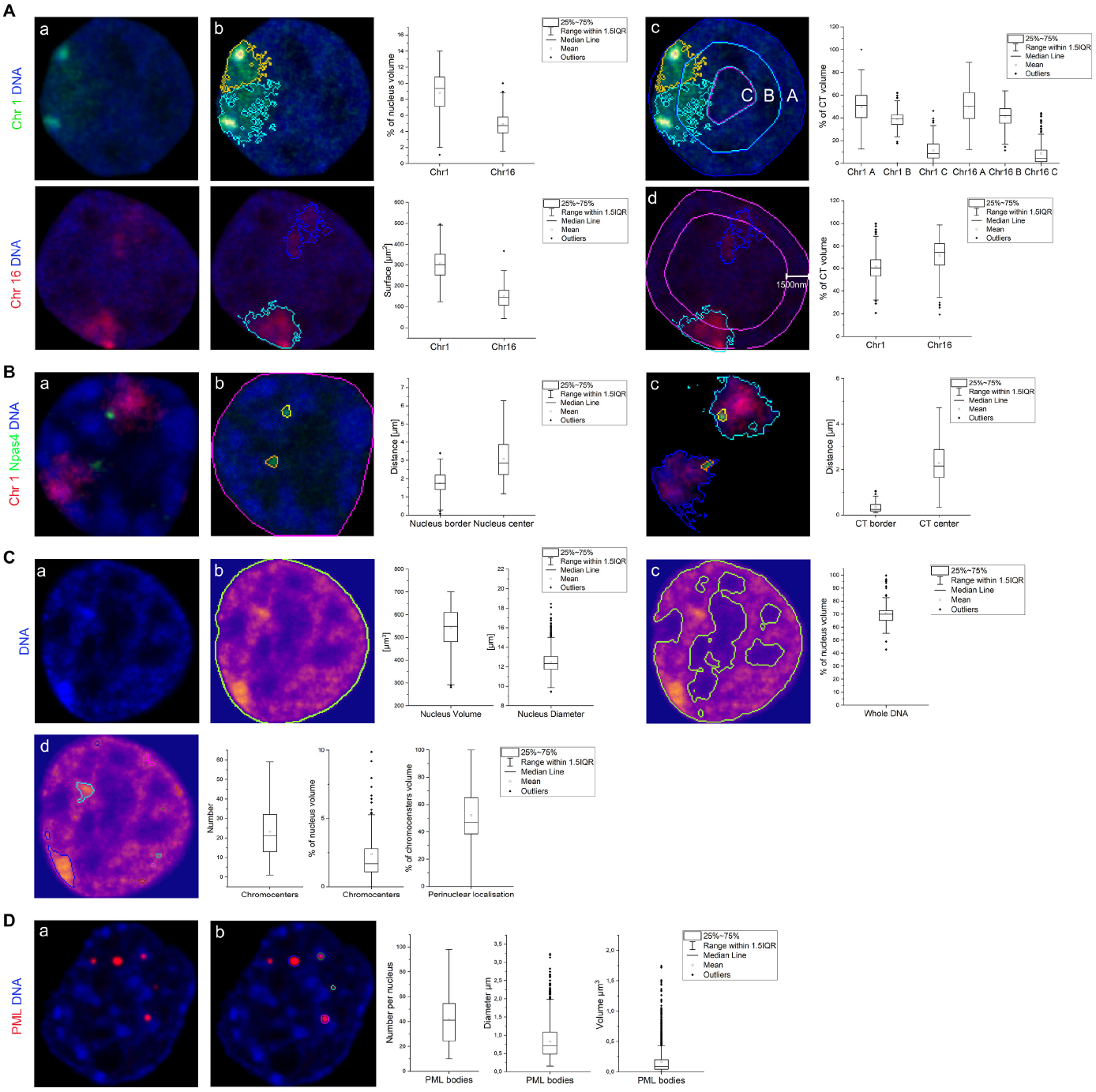
Use cases showing examples of PartSeg analysis performed on 3D confocal images of rat hippocampal neurons and mouse neuroblastoma cell line. A. Distribution of chromosome 1 and 16 territories (CTs) in a neuronal nucleus. a) Confocal picture of a single, segmented nucleus with CT 1 (upper panel, green) and CT 16 (lower panel, red) visualised by FISH. b) Quantification of volume and surface of CT 1 (segmented with MSO algorithm) and CT 16 (segmented with manual threshold). c) Radial distribution of CTs 1 and 16 volume within a neuronal nucleus. Concentric spheres depicted for chromosome 1 as A, B and C have equal radius. d) Perinuclear distribution of CT 1 and 16. CT volume located within 1500 nm from the nucleus boundary (depicted for chromosome 16) was measured for each chromosome.B. Distribution of Npas4 gene within a neuronal nucleus and chromosome 1 CT. a) Confocal picture of a single, segmented nucleus with chromosome 1 (depicted in red) and Npas4 gene (depicted in green) visualised by FISH. b) Segmentation of Npas4 gene and its localization in a neuronal nucleus. c) Segmentation of Npas 4 gene and chromosome 1 CT, graphs show Npas4 localisation relative to CT. C. Neuronal nucleus and chromatin analysis. a) DNA staining of a neuronal nucleus. b) Segmentation of neuronal nucleus based on DNA staining. Quantification of nucle1a7r diameter and volume. c) Quantification of average DNA volume based on segmentation using Otsu automated threshold. d) Quantification of chromocenters based on segmentation using Rényi Entropy automated threshold, graphs show chromocenters number, relative volume and percentage of chromocenters volume localised within 800 nm from the nuclear border.D. Quantification of PML bodies. a) Confocal picture of a single, segmented nucleus of a mouse neuroblastoma cell line with PML bodies depicted in red. b) Segmentation of PML bodies and quantification of their number, diameter and volume.

Automatic segmentation of nuclear structures poses many challenges because of confinement boundaries imposed b y the nucleus, dense packing and significant background noise, resulting in signal overlap. Optimal segmentation of boundaries via intensity-threshold based methods is difficult, if not impossible, especially for conjoined structures. Yet, segmentation of ROI is essential for expert evaluation and serves as basis for calculating numerical descriptors of data.

Because of this, there is no possibility of creating a parameterless, universal algorithm for processing data from innovative experiments. Rather a flexible methodology for tuning data processing methods to current datasets is preferable. We take this task one step further by facilitating the process for users without programmatic knowledge.

In this paper, we present PartSeg, which offers a novel segmentation algorithm for a high-throughput imaging data and is specially optimised for analysis of cell nucleus. PartSeg relies on 3D segmentation of regions of interest (ROI) and subsequent analysis over a large number of morphological features of the segmented structures. It is equipped with an easy-to-use graphical user interface (GUI) and can alternatively be utilised as a Python library in composite data processing pipelines. Therefore, it meets the expectations of both biologists and bioinformaticians. Moreover, PartSeg allows for batch processing of datasets coming from different sources e.g. 2D/3D images from light or electron microscopy, which can be managed without the support of a programmer by the use of rapid prototyping on sample data.

The main design goal of PartSeg was to make it simple to learn for new users. To achieve this, we determined typical use cases for users interested in studying cell nuclei (see Subsection 2.1) and planned the GUI so that swift proficiency in usage of the toolkit can be quickly achieved. It is consistent, ergonomic and does not introduce unnecessary notions or mechanisms. For example, compatibility is maintained by supporting a diverse set of 3D image formats stored in the filesystem enabling interoperability with all popular platforms. This way no importing, or preprocessing is needed and user can focus on the tasks at hand, rather than the software’s inner workings. This follows Human Computer Interaction studies [5], which show that users expect systems to work and will chose those that are easier to use.

The current revision of the PartSeg interface includes support for 2D and 3D multichannel data. It consist of two separate tools named *Mask Segmentation* and *ROI Analysis*. The former is intended for extraction of objects of interest from the data set and can store the basic information on those objects in a separate mask file. The latter can be used for the measurements of morphological parameters of specific structures within the extracted objects.

Finally, some ergonomics improvements are incorporated following examination of typical usage patterns. For example, segmentation parameters are automatically saved during and in between sessions, and specific segmentation profiles can be saved by users. The segmented a rea is graphically presented on top of the data which simplifies visual validation of utilised parameters. PartSeg is also equipped with a synchronised view, where two windows are used simultaneously, permitting the user to compare the current segmentation with multichannel raw data or with other segmentation computed with different parameters. Last but not least, parallel batch processing is available, with the number of processes adjusted by the user depending on available system resources.

## 2 Implementation

In order to optimise Partseg GUI we have identified typical steps of nucleus analysis workflow by examining several real life use cases. In subsection 3.2 we provide four examples of such workflow.

### 2.1 Typical steps of nucleus analysis workflows

The steps are performed manually on a small subset of the data. This confirms the selection of the method and that the parameters are adjusted appropriately before processing of the whole dataset.

#### Preselection of nuclei in 3D images

In order to quantitatively analyse structures within cell nucleus, it is necessary to segment individual nuclei from the 3D image. Nuclei can be segmented based on any nuclear staining and thresholding method. However, in the case of heterogeneous samples with diverse staining intensities, one threshold is not enough to properly segment all nuclei. The possibility of limiting the selection of nuclei from the population for analysis provides opportunity to study cells in particular cell cycle phase or those of a specific type e.g. single cancerous cells embedded in a normal tissue.

#### Selection and verification of parameters for extraction of complex ROI inside the nucleus’ territory

Next, segmentation of ROI within nuclei is necessary in order to obtain numerical values for nuclear assemblies. Extraction of complex structures from 3D images usually requires testing of several algorithms in order to select optimal parameters. Visualisation of performed segmentation on top of raw data allows the user to select methods and settings facilitating the extraction process. As was mentioned before, nuclear space is very crowded. For example pairs of chromosome territories (CTs) are often localised close to each other, which makes segmentation of a single copy very difficult even for an experienced eye. This step is tuned on a sample subset of nuclei.

#### Measurement set

A choice of numerical parameters (measurements) needs to be made. Values for those parameters are calculated for each image as a result of manual or batch segmentation processing and presented to the user in the form of a spreadsheet. The user then performs some simple computations and plots the results. Results should be based on the initial picture resolution and expressed in physical units.

#### Batch processing

When single image analysis is considered satisfactory, it can be repeated by executing batch processing on all input images to collect data for statistical analysis. Results are shown for each individual nucleus and structure. This allows conclusions to be drawn for the entire cell population based on data acquired from individual nuclei simplifying the subsequent tasks of categorising nuclei or structures and performing statistical analysis. Comparing, aforementioned biochemical techniques show an averaged outcome from large populations of cells, which does not necessarily reflect the biological heterogeneity of individual components.

Although several general and mature bioimaging tools like ImageJ [6], Icy [7], CellProfiler [8] and ImagePy [9] exist, none of these provide support for all of the aforementioned steps nor do they possess an interface optimised for the workflow as a whole. A detailed comparison with other systems is provided in Section 3.1.

### 2.2 Algorithms for ROI Analysis

The main feature of PartSeg is ROI analysis. ROI is a set of voxels, which can be distinguished and measured. In PartSeg there are several provided algorithms for that, which we list below, and more can be added as plugins. Here, we categorise the algorithms into two groups. The first contains algorithms, which are designed for ROI extraction from an initial 3D image, outputting a collection of ROI. The second contains algorithms for measuring various ROI features. These algorithms take a collection of ROIs as an input and use it to compute numerical or aggregate features.

#### 2.2.1 ROI extraction

The canonical group of PartSeg algorithms designed for finding collections of ROI rely on threshold segmentation. These include:

- Thresholding — ROI is defined by thresholding followed by identification of connected components and minimum size filtering. The upper or lower threshold level can be set manually by the user or calculated with common methods such as Otsu, Shanbhag, etc.
- Range thresholding — allows user to set specific threshold range marking ROI, which can be useful e.g. to eliminate background staining.
- Thresholding with watershed — ROI is found by two stage thresholding. First core objects are identified as in *Thresholding*, after which a second thresholding marks the whole area of interest. Voxels from this area are found using selected watershed-like methods initiated from core objects.
- Multiscale opening — is a type of watershed-like thresholding that is unique to PartSeg. Further details are given in Subsection 2.3.
- Multiple threshold Otsu — is a generalisation of the histogram-based Otsu method (see [10]) which identifies multiple types of ROIs using set of thresholds.

#### 2.2.2 Plugins

PartSeg has plugin system which allows to expand it with additional features. This is convenient if one wants to add experiment specific computational methods and include tailored dependencies that need to be selected to match specific computer configuration. As an example we have provided plugins incorporating deep learning algorithms. It is well recognised that Deep Learning gives very good results, when applied to the nucleus segmentation. There are many published models (both networks topologies, and their software implementations), for example Stardist [11, 12] and Cellpose [13]. Typically such highly-optimised statistical methodologies have specific configurations that depend on processor type, graphic card and installed drivers, hence we implement them as plugins. Our plugins for these two models are available at: https://pypi.org/project/PartSeg-stardist/ and https://pypi.org/project/PartSeg-cellpose/.

ROI selection can be performed multiple times in an incremental manner. After determining the first collection, selected ROI can be converted to a mask and used as an input for the next level of extraction. When the final collection is obtained, its numerical description can be determined using algorithms described in the next section.

#### 2.2.3 Measuring of ROI features

Measurements can be performed on an area defined with *ROI*, collection of *ROI* or a *Mask*. Several common measurement methods like *Volume* or *Diameter* are available which require no explanation.

PartSeg also provides more sophisticated methods of ROI measurements on Mask. They can, for example, be used to measure gene positions in the nucleus or to measure relative difference of concentrations of proteins within regions of the nucleus. These methods include:

- Border rim — it allows to measure the total volume or pixel brightness of the selected ROI, which is located within a given distance from the border of mask. An example application would be to identify gene or the portion of chromosome territory positioned in close proximity to nuclear rim (see Figure 1 A.d. and [14])
- Mask distance splitting — splits the mask into concentric regions of increasing distance from the mask centre, which can be of equal radius or equal volume. It allows to measure volume and pixel brightness of ROI found within the designated regions. For example it shows radial position of CTs within nucleus (see Figure 1 A.c. and [15]).
- Mask-ROI distance — distance from ROI to mask is calculated based on their *mass centre* (taking brightness into account), *geometrical centre*, or *border distance*.

An example of application would be to identify gene positioning within the nucleus (see Figure 1 A.d. and [14])

Some other noteworthy measurement types include:

- Moment — One of possible measurements of mass (pixel brightness) distribution inside ROI. It allows us to determine if the structural mass is concentrated or distributed evenly. The formula is 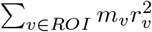, where *m*_*v*_ is the brightness of a voxel and *r*_*v*_ is its distance from the ROI’s centre of mass. The interpretation is similar to the classical moment of inertia. This measure assume that voxel brightness is one to one correlated with object density.
- First/Second/Third principal axis — Aligns ROI using weighted PCA, then calculates ROI length along the corresponding axis. The weights correspond to voxel brightness and their position vector is in relation to the ROI’s centre of mass. This measurement can be used to determine basic shape of ROI.

### 2.3 Multiscale Opening

Often in the analysis of imaging data the distance between two ROIs is smaller than the angular resolution of imaging method used. Historically, the first approach to separate such ROIs was the watershed [16] method. However if an object has a diverse morphology i.e. exhibits regions of higher brightness, classical watershed will incorrectly split such an object into multiple ROIs. To overcome this problem, the watershed transform was developed [17]. Unfortunately watershed transform works best for objects, which are spherical in a chosen metric. Because many nuclear domains are densely packed and non-spherical, it was important to develop methods that are capable of reliable segmentation. Therefore we implemented a novel Multiscale Opening [18]. It takes into account both the change in brightness and the physical distance along the joining path. The main difference between other watershed like algorithms and Multiscale Opening is its iterative voxel labelling. Voxels can be labelled only if they are closer to any object than to the background. This approach produces better results for stretched and non-symmetrical objects. An example is presented in Figure 3.

### 2.4 Tutorials

PartSeg can be used as a standalone program or as a Python library for example, as part of a larger pipeline. We provide two tutorials, which show how to use PartSeg from the biologists perspective with the Graphical User Interface (GUI) and from the bioinformaticians perspective as a library.

#### 2.4.1 Using PartSeg GUI

https://github.com/4DNucleome/PartSeg/blob/master/tutorials/tutorial-chromosome-1/tutorial-chromosome1_16.md. In this tutorial we present how PartSeg can be used to segment nuclei from 3D confocal images and subsequently analyse several parameters of chromosomal territories in chromosomes 1 and 16. This use case can be broken down into three parts.

First, the segmentation of nuclei is performed using the DNA signal. Segmented nuclei are cut from the original pictures and the mask files containing segmentation parameters are created.

Second, in order to quantify features of chromosome 1 territories (CT1), segmentation of its specific signal is carried out. The parameters for segmentation are adjusted to accurately cover the chromosome’s staining. Next, the method for measuring several morphological parameters of CT1 is presented. These parameters are calculated based on a fixed threshold value. Additionally, the volume ratio of CT1 and its nucleus is calculated.

Finally, the previously established settings profile is used in batch mode to measure features of the nuclei and CT1s. After repeating the process for chromosome 16, a comparison of size between chromosome 1 and 16 can be done in a fully automated way (see 1) A. In Figure 2 we utilise the PartSeg GUI to perform steps from the typical workflow in Section 2.1.

**Figure 2:**
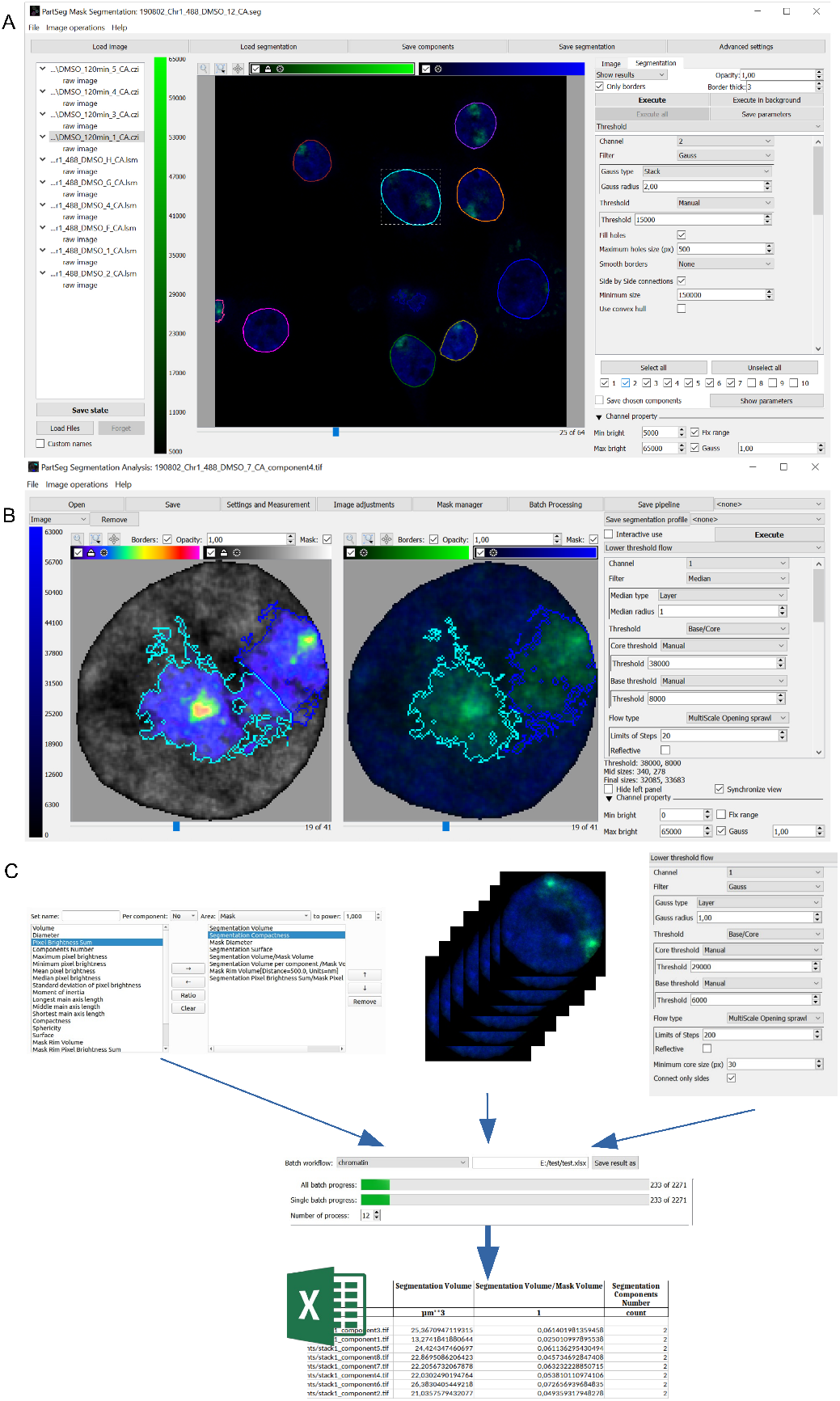
GUI overview **A** GUI for nucleus segmentation and preselection, **B** Main GUI for analysis of single cases and preparation of parameters for batch processing, **C** In batch mode, the choice of the measurement set results in a spreadsheet with corresponding columns and with results for all images.

**Figure 3:**
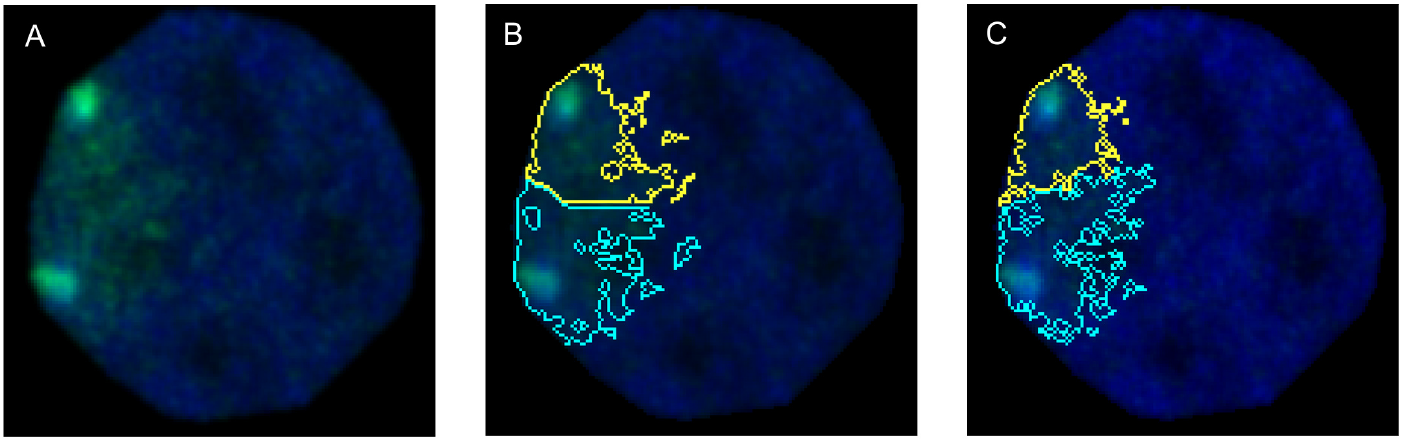
Watershed and Multiscale Opening Comparison. Difference between object separation with Multiscale Opening and Watershed transform. A) Confocal picture of a single, segmented nucleus with FISH signal from a chromosome 1 probe (depicted in green). B) Segmentation of chromosome 1 signal using the Watershed algorithm. C) Segmentation of chromosome 1 signal using the Multiscale Opening algorithm.

**Figure 4:**
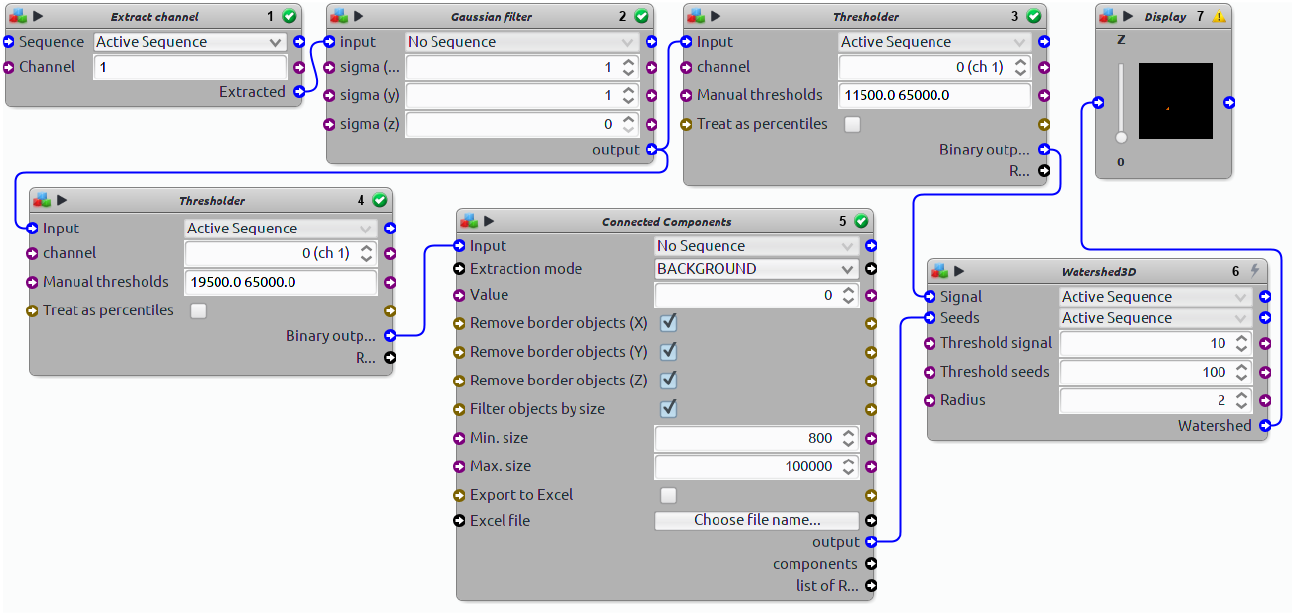
Sample protocol from Icy. It contains approximation, using blocks from standard installation of *Thresholding with watershed* ROI extraction method from 2.2.1.

#### 2.4.2 Using PartSeg as a Python library

https://github.com/4DNucleome/PartSeg/blob/master/tutorials/tutorial_neuron_types/Neuron_types_example.ipynb In this tutorial we present how PartSeg components can be used as a Python library. Images of cells immunostained for markers Prox1 and CamKII and counterstained with DNA dye were acquired with confocal microscopy. Initially, segmentation of hippocampal neuron nuclei is shown. Next, a combination of PartSeg components and custom code is presented in order to assign segmented nuclei to 4 different classes based on the aforementioned markers. Finally, segmentation of chromatin based on DNA staining is performed to obtain a set of measurements. In the final step, Matplotlib plots of the obtained data are generated. Tutorial shows how to create aforementioned pipeline using API method for segmentation, or simply load it from a file exported in PartSeg GUI.

API documentation of PartSeg is available at https://partseg.readthedocs.io/en/stable/.

## 3 Discussion

### 3.1 Comparison to existing tools

There exist several general tools for image processing and analysis like ImageJ [6], Icy [7], CellProfiler [8]. These are robust apps that have been developed over years and contain a wealth of options and plugins for many applications. For inexperienced users such a multiplicity is difficult to overcome and the general nature of these tools does not enforce a workflow adapted to nucleus analysis.

ImageJ’s, Icy’s and ImagePy’s main workflow revolves around atomic operations like thresholding or filtering. In those tools presentation of processing results in the context of input data requires many additional steps. For example Icy allows for the creation of a pipeline in a graphical setting, but does not feature a simplified view or e xport of key parameters 4. This type of work resembles graphical programming, where user needs to think in the context of atomic operations and not a semantic meaning. For now such editor is unavailable for ImageJ and ImagePy, both offer only scripting, which demands basic programming knowledge.

CellProfiler provides a tool for pipeline creation and execution, however it does have an exploration mode. Although it gives a possibility to preview intermediate results, the implemented viewer was not designed for 3D data, therefore it is restricted to only a few layers automatically selected by the tool. The user is expected to define a multi-step computational pipeline himself, this often proves too difficult a task to overcome without programmatic experience and results in an interface less ergonomic in this respect to one offered by PartSeg.

We have taken an alternative approach, where the GUI is simple, compact and provides easy exploration of various algorithms and their wide range of parameters with immediate visualisation of results. The GUI is organised around the workflow, which we have defined with our users (see subsection 2). Even though our central focus is on analysis of cell nuclei, Partseg is general enough to cover various other use cases. The modular structure of PartSeg allows saving of ROI segmentation parameter settings at many levels of complexity: from simple profiles, through pipelines, to projects containing entire segmentation and imaging data. This enables easy collaboration and facilitates repeatable and verifiable research as the entire segmentation can be published as supplementary material. On the other hand power users can integrate PartSeg in their analysis pipelines with the use of Python API that we provide.

From the two other tools dedicated specifically to analysis of cell nucleus, NEMO [19] and TANGO [20] only the latter still can be obtained. TANGO, reviewed in [21], requires installation of external dependencies to work with a full efficiency. Such installation might be challenging for a typical user and cannot be done without administrative privileges. Moreover TANGO’s GUI seems to be highly unintuitive. It seems that all aforementioned tools are better suited for bioinformaticians, than wet lab scientist. Yet, it is a wet lab scientist, who the best understands whole experimental setup and visual outcome and can asses if ROI is segmented properly. The philosophy behind PartSeg is to provide GUI equipped in high level operations with semantic descriptions, as well as Python library with API. Implementation details are hidden from the user, but at the same time they are empowered with multichannel 3D viewer, capable of showing ROI in the context of input data and recording the analysis parameters for reuse on other datasets. Users with more advanced programming knowledge can use the Python API and easily combine PartSeg with other data analysis libraries in their processing pipelines.

### 3.2 Examples of PartSeg Application on Real Data

This section contains four examples of the workflow a forementioned in subsection 2 applied to 3D confocal images of rat hippocampal neurons cultured *in vitro* (Fig. 1 A-C) and mouse neuroblastoma cell line (Fig. 1 D). Graphics and values shown on Fig. 1 were obtained using PartSeg.

First, we analysed the surface, relative volume and distribution of territories of chromosomes 1 and 16 in nuclei of rat hippocampal neurons (Fig. 1 A). The chromosomes were stained using FISH with chromosome 1 and 16 paint probes. The measured volume of both chromosomes territories (CT) roughly correlates with their size in Mbp-267.9 Mbp for chromosome 1 and 90.2 Mbp for chromosome 16, which is 10 and 3,5 % of the whole genome accordingly. Some discrepancies are expected due to varying distributions of heterochromatin and euchromatin in both chromosomes, as well as conditions of hybridisation, which require DNA heat denaturation. PartSeg allows us to analyse radial distribution of segmented structures within the nucleus (Fig. 1. A, c), together with their proximity to the nuclear border (Fig. 1. A, d and B, b). We checked localisation of both CTs in 3D nuclear space. Both CTs were in close proximity to the nuclear periphery (Fig. 1 A, c and d) and on average most of the CT volume was located within 1500 nm from the nucleus boundary (50% of the volume of CT1 and 70% of CT16). It was shown that chromatin of mouse and human cells show presence of lamina associated domains (LADs) distributed along all chromosomes, which cover around one-third of the whole genome, therefore all chromosomes are in contact with nuclear lamina located at the nuclear border [22]. However the spatial organisation of CTs is flexible, permitting many local and long-range contacts of genes and regulatory elements, which influence their function [23].

Data suggest that radial positioning of genes often correlates with a transcriptional state, with actively transcribed genes located in the interior and silent genes at the periphery of the nucleus. Also, the gene position within CT reflects its state of expression, where active genes tend to localise the CT boundary [24]. Therefore, we have analysed the distribution of the Npas4 gene in rat hippocampal neurons subjected to sequential FISH with chromosome 1 paint probe and Npas4 probe (Fig. 1 B). Npas4 is a transcription factor involved in structural and functional plasticity, actively transcribed in neurons [25]. In mature rat hippocampal neurons Npas4 gene was located close to the border of chromosome 1 (Fig1. B, c) at the same time remaining relatively far from the nuclear periphery (Fig. 1 B, b), which is in agreement with the aforementioned observations for active genes.

Next, we looked at nuclei and chromatin of mature rat hippocampal neurons (Fig1. C). For 750 nuclei we have determined an average volume of 556 *μ*m^3^ and a diameter of 12,5 *μ*m (Fig1. C, b). DNA stained with Hoechst dye, occupied on average 70% of the nucleus volume (Fig1. C, c). We have also calculated the average number and volume of chromocenters, which contain highly condensed and constitutively silenced, pericentromeric chromatin (Fig1.C, d). We found an average number of 20 chromocenters per nucleus, which occupied around 2% of the nuclear volume, with 50% of them localised within 800 nm of the nuclear border. Quantification of e.g. chromatin volume is interesting as its condensation is a dynamic process dependent on many physiological and environmental factors [26]. Chromatin compaction changes accompany cell differentiation, cell division, cell death, senescence, ischaemia and oxidative stress as well as constitutive expression silencing [27].

As the last example we have calculated the number, diameter and volume of PML bodies in mouse neuroblastoma cell lines (Fig. 1 D). PML (promyelocytic leukaemia) bodies are matrix associated, spherical, nuclear bodies, with a diameter of 0.1–1.0 *μ*m, which can be found in most cell lines and many tissues. Nuclei of asynchronous neuroblastoma cell lines had on average 40 PML bodies per nucleus, with a diameter of 0,8 *μ*m and volume of 0,17 *μ*m^3^. It was shown that size and number of PML bodies heavily depends on cell cycle phase, cell type and stage of differentiation. Changes of the number and volume of these nuclear domains accompany induction of stress, senescence and tumorgenesis [28].

Figure 1 presents some examples of nuclei and nuclear domains, like chromosome territories, genes, PML bodies, chromatin and chromocenters, which can be quantitatively analyzed in PartSeg. However, PartSeg can easily be adapted to quantify other cellular structures like mitochondria or Golgi apparatus.

## 4 Conclusions

In this paper, we have presented PartSeg which is a tool for precise and verifiable segmentation and reconstruction of 3D microscopy images. PartSeg is optimised for cell nucleus analysis and offers multi-scale segmentation algorithms best-suited for this task. PartSeg can also be used for bulk processing of multiple images and its components can be reused programmatically in other systems or computational experiments.

Furthermore, simple storing of the whole segmentation process with settings and data, empowers cooperation and independent verification of results in the spirit of data provenance [29], and open and verifiable science.

## 5 Declarations

### 5.1 Ethics approval and consent to participate

Not applicable.

### 5.2 Consent for publication

Not applicable.

### 5.3 Availability of data and materials

**Project name:** PartSeg

**Project home page:** https://4dnucleome.cent.uw.edu.pl/PartSeg/, https://github.com/4DNucleome/PartSeg

**Operating system(s):** Platform independent

**Programming language:** Python, C++

**Other requirements:** Python 3.6+

**License:** BSD (with GPL dependencies)

**Tutorials and documentation**: http://4dnucleome.cent.uw.edu.pl/PartSeg/.

## 5.4 List of abbreviations

3-C: Chromosome Conformation Capture - method of chromatin interaction study
API: Application Programming Interface
BSD: Berkeley Software Distribution Licenses https://www.gnu.org/licenses/license-list.en.html#ModifiedBSD
CTs: Chromosome territories
GPL: GNU General Public License https://www.gnu.org/licenses/license-list.en.html#GNUGPL
GUI: Graphical User Interface
PML: Promyelocytic leukemia proteins
ROI: Regions of Interest
Mbp: Mega base pairs

## 5.7 Authors’ contributions

GB and DP proposed the project and the software concept; GB performed the code development; DP, AM, JS provided the guidance and the project management; GB, AM, JS and DP were involved in software testing; PT, AG, YY and AM performed the biological experiments; SB and ND developed the multiscale opening (MSO) segmentation algorithm; GB implemented the MSO in PartSeg; JS, AM, PT, DP, GB were responsible for the writing of the manuscript. All authors read and approved the final manuscript.

## 5.8 Acknowledgements

This paper is dedicated as a memorial to Professor Grzegorz Wilczyński, who suddenly passed away on 13 July 2020. Grzegorz has contributed to the idea and initial development of PartSeg.

## 5.9 Authors’ information

## References

[1] Peddie, C.J., Collinson, L.M.: Exploring the third dimension: volume electron microscopy comes of age. Micron 61, 9–19 (2014)

[2] Pegoraro, G., Misteli, T.: High-throughput imaging for the discovery of cellular mechanisms of disease. Trends in Genetics 33(9), 604–615 (2017). doi:10.1016/j.tig.2017.06.005

[3] Woodcock, C.L.: Chromatin architecture. Current Opinion in Structural Biology 16(2), 213–220 (2006). doi:10.1016/j.sbi.2006.02.005

[4] Fraser, P., Bickmore, W.: Nuclear organization of the genome and the potential for gene regulation. Nature 447(7143), 413–417 (2007). doi:10.1038/nature05916

[5] Dix, A., Finlay, J., Abowd, G.D., Beale, R.: Human Computer Interaction, 3rd edn. Pearson Prentice Hall, Harlow, England (2003)

[6] Rueden, C.T., Schindelin, J., Hiner, M.C., DeZonia, B.E., Walter, A.E., Arena, E.T., Eliceiri, K.W.: ImageJ2: ImageJ for the next generation of scientific image data. BMC Bioinformatics 18(1), 529 (2017). doi:10.1186/s12859-017-1934-z

[7] De Chaumont, F., Dallongeville, S., Chenouard, N., Hervé, N., Pop, S., Provoost, T., Meas-Yedid, V., Pankajakshan, P., Lecomte, T., Le Montagner, Y., Lagache, T., Dufour, A., Olivo-Marin, J.C.: Icy: An open bioimage informatics platform for extended reproducible research. Nature Methods 9(7), 690–696 (2012). doi:10.1038/nmeth.2075

[8] Carpenter, A.E., Jones, T.R., Lamprecht, M.R., Clarke, C., Kang, I.H., Friman, O., Guertin, D.A., Chang, J.H., Lindquist, R.A., Moffat, J., Golland, P., Sabatini, D.M.: CellProfiler: image analysis software for identifying and quantifying cell phenotypes. Genome Biology 7(10), 100 (2006). doi:10.1186/gb-2006-7-10-r100

[9] Wang, A., Yan, X., Wei, Z.: ImagePy: an open-source, Python-based and platform-independent software package for bioimage analysis. Bioinformatics 34(18), 3238–3240 (2018). doi:10.1093/bioinformatics/bty313

[10] Liao, P.-S., Chen, T.-S., Chung, P.-C.: A fast algorithm for multilevel thresholding. J. Inf. Sci. Eng. 17, 713–727 (2001)

[11] Schmidt, U., Weigert, M., Broaddus, C., Myers, G.: Cell detection with star-convex polygons. In: Medical Image Computing and Computer Assisted Intervention - MICCAI 2018 - 21st International Conference, Granada, Spain, September 16-20, 2018, Proceedings, Part II, pp. 265–273 (2018). doi:10.1007/978-3-030-00934-2_30

[12] Weigert, M., Schmidt, U., Haase, R., Sugawara, K., Myers, G.: Star-convex polyhedra for 3d object detection and segmentation in microscopy. In: The IEEE Winter Conference on Applications of Computer Vision (WACV) (2020)

[13] Stringer, C., Michaelos, M., Pachitariu, M.: Cellpose: a generalist algorithm for cellular segmentation. bioRxiv (2020). doi:10.1101/2020.02.02.931238. https://www.biorxiv.org/content/early/2020/02/03/2020.02.02.931238.full.pdf

[14] Walczak, A., Szczepankiewicz, A.A., Ruszczycki, B., Magalska, A., Zamlynska, K., Dzwonek, J., Wilczek, E., Zybura-Broda, K., Rylski, M., Malinowska, M., et al.: Novel higher-order epigenetic regulation of the bdnf gene upon seizures. Journal of Neuroscience 33(6), 2507–2511 (2013)

[15] Tanabe, H., Müller, S., Neusser, M., von Hase, J., Calcagno, E., Cremer, M., Solovei, I., Cremer, C., Cremer, T.: Evolutionary conservation of chromosome territory arrangements in cell nuclei from higher primates. Proceedings of the National Academy of Sciences 99(7), 4424–4429 (2002). doi:10.1073/pnas.072618599. https://www.pnas.org/content/99/7/4424.full.pdf

[16] Soille, P., Vincent, L.M.: Determining watersheds in digital pictures via flooding simulations. In: Kunt, M. (ed.) Visual Communications and Image Processing ’90: Fifth in a Series. SPIE, ??? (1990). doi:10.1117/12.24211. https://doi.org/10.1117/12.24211

[17] B.T.M., R.J., Arnold, M.: The watershed transform: Definitions, algorithms and parallelization strategies. Fundamenta Informaticae 41(1, 2), 187–228 (2000). doi:10.3233/FI-2000-411207

[18] Saha, P.K., Basu, S., Hoffman, E.A.: Multiscale opening of conjoined fuzzy objects: Theory and applications. IEEE Transactions on Fuzzy Systems 24(5), 1121–1133 (2016)

[19] Iannuccelli, E., Mompart, F., Gellin, J., Lahbib-Mansais, Y., Yerle, M., Boudier, T.: Nemo: a tool for analyzing gene and chromosome territory distributions from 3d-fish experiments. Bioinformatics 26(5), 696–697 (2010). doi:10.1093/bioinformatics/btq013./oup/backfile/content_public/journal/bioinformatics/26/5/10.1093/bioinformatics/btq013/2/btq013.pdf

[20] Ollion, J., Cochennec, J., Loll, F., Escudé, C., Boudier, T.: Tango: a generic tool for high-throughput 3d image analysis for studying nuclear organization. Bioinformatics 29(14), 1840–1841 (2013). doi:10.1093/bioinformatics/btt276./oup/backfile/content_-public/journal/bioinformatics/29/14/10.1093_bioinformatics_-btt276/2/btt276.pdf

[21] Chessel, A.: An overview of data science uses in bioimage informatics. Methods 115, 110–118 (2017). doi:10.1016/j.ymeth.2016.12.014. Image Processing for Biologists

[22] van Steensel, B., Belmont, A.S.: Lamina-associated domains: Links with chromosome architecture, heterochromatin, and gene repression. Cell 169(5), 780–791 (2017). doi:10.1016/j.cell.2017.04.022

[23] Bickmore, W.A.: The spatial organization of the human genome. Annual Review of Genomics and Human Genetics 14(1), 67–84 (2013). doi:10.1146/annurev-genom-091212-153515. PMID: 23875797. https://doi.org/10.1146/annurev-genom-091212-153515

[24] Tang, Z., Luo, O.J., Li, X., Zheng, M., Zhu, J.J., Szalaj, P., Trzaskoma, P., Magalska, A., Wlodarczyk, J., Ruszczycki, B., et al.: CTCF-mediated human 3D genome architecture reveals chromatin topology for transcription. Cell 163(7), 1611–1627 (2015)

[25] Sun, X., Lin, Y.: Npas4: Linking neuronal activity to memory. Trends in Neurosciences 39(4), 264–275 (2016). doi:10.1016/j.tins.2016.02.003

[26] Ravi, M., Ramanathan, S., Krishna, K.: Factors, mechanisms and implications of chromatin condensation and chromosomal structural maintenance through the cell cycle. Journal of Cellular Physiology 235(2), 758–775 (2020). doi:10.1002/jcp.29038. https://onlinelibrary.wiley.com/doi/pdf/10.1002/jcp.29038

[27] Illner, D., Zinner, R., Handtke, V., Rouquette, J., Strickfaden, H., Lanctôt, C., Conrad, M., Seiler, A., Imhof, A., Cremer, T., Cremer, M.: Remodeling of nuclear architecture by the thiodioxoxpiperazine metabolite chaetocin. Experimental Cell Research 316(10), 1662–1680 (2010). doi:10.1016/j.yexcr.2010.03.008

[28] Lallemand-Breitenbach, V., de Thé, H.: Pml nuclear bodies: from architecture to function. Current Opinion in Cell Biology 52, 154–161 (2018). doi:10.1016/j.ceb.2018.03.011. Cell Nucleus

[29] Cuevas, V., Dey, S., Köhler, S., Riddle, S., Ludäscher, B.: Scientific workflows and provenance: Introduction and research opportunities. Datenbank-Spektrum 12(2013). doi:10.1007/s13222-012-0100-z

